# Simultaneous Epigenetic Perturbation and Genome Imaging Reveal Distinct Roles of H3K9me3 in Chromatin Architecture and Transcription

**DOI:** 10.1101/2020.07.15.204719

**Authors:** Ying Feng, Yao Wang, Xiangnan Wang, Xiaohui He, Chen Yang, Ardalan Naseri, Thoru Pederson, Jing Zheng, Shaojie Zhang, Xiao Xiao, Wei Xie, Hanhui Ma

## Abstract

Despite the long-observed correlation between H3K9me3, chromatin architecture and transcriptional repression, how H3K9me3 regulates genome higher-order organization and transcriptional activity in living cells remains unclear. Here we develop EpiGo (<u>E</u>pigenetic <u>p</u>erturbation <u>i</u>nduced <u>G</u>enome <u>o</u>rganization)-KRAB to introduce H3K9me3 at hundreds of loci spanning megabases on human chromosome 19 and simultaneously track genome organization. EpiGo-KRAB is sufficient to induce *de novo* heterochromatin-like domain formation, which requires SETDB1, a methyltransferase of H3K9me3. Unexpectedly, EpiGo-KRAB induced heterochromatin-like domain does not result in widespread gene repression except a small set of genes with concurrent loss of H3K4me3 and H3K27ac. Ectopic H3K9me3 appears to spread in inactive regions but is largely restricted to transcriptional initiation sites in active regions. Finally, Hi-C analysis showed that EpiGo-KRAB induced to reshape existing compartments. These results reveal the role of H3K9me3 in genome organization could be partially separated from its function in gene repression.

## Background

Human genome is organized in a hierarchy manner from kilobase to megabase scales such as nucleosome, loops, topologically associated domains (TADs) and A/B compartments [1–4]. It has been proposed that the loop extrusion drives TAD formation [5]. On the other hand, liquidliquid phase separation is suggested to mediate genome compartmentalization [5, 6]. For example, heterochromatin protein HP1α undergoes liquid-liquid demixing suggesting a role of phase separation in heterochromatin domain formation [7–9]. Heterochromatin drives compartmentalization in the inverted nuclei of rods in nocturnal mammals [10]. The cosegregated compartments often share similar chromatin states such as histone marks [11]. Despite the widely observed correlations [11], how the epigenetic modifications regulates genome organization, particularly in living cells, remains unclear. Direct visualization of chromatin structures in cell nucleus is still challenging. OligoSTORM, MERFISH, Hi-M or ORCA has been applied to trace DNA folding [12–18] in fixed cells. On the other hand, CRISPR-based imaging provides a versatile and powerful tool to track chromatin topology in live cells in real time [19, 20].

Here we develop a CRISPR-based EpiGo-KRAB system to investigate the effect of ectopic H3K9me3 on genome organization and transcription in living cells. We show EpiGo-KRAB of a large chromatin domain induces *de novo* heterochromatin-like domain formation and reshapes local compartmentalization. Surprisingly EpiGo-KRAB induced heterochromatin-like domain does not result in global gene repression. We believe that this system should be applicable for other epigenetic perturbations.

## Results

### Establishment of EpiGo-KRAB

To investigate how H3K9me3 regulates genome architecture and gene expression in living cells, here we developed a CRISPR-based system, namely EpiGo (<u>E</u>pigenetic <u>p</u>erturbation <u>i</u>nduced <u>G</u>enome <u>o</u>rganization)-KRAB (**Fig. 1A)**. The EpiGo-KRAB system allows to epigenetic manipulation of defined regions and visualizing the subsequent spatiotemporal dynamics of these loci. First, we utilized dCas9-KRAB, which deposits loci-specific epigenetic modifications [21, 22], and fluorescent guide RNAs from the CRISPRainbow system (sgRNA-2XPP7 and PCP-GFP) for DNA visualization [23]. We intended to alter the epigenetic states from kilobases to megabase scales, to study the role of epigenetic modification in genome organization at different scales. By mining the chromosome-specific repeats across megabases of human genome, we found a repeat class which consists of 836 copies of CRISPR target sites spanning ~17 megabases at the q-arm of chromosome 19, and we dubbed it C19Q as EpiGo-KRAB targets for the following studies (**Fig. 1A and Table S1**). C19Q can be visualized by co-expression of dCas9-KRAB, sgRNA-2XPP7 and PCP-GFP. We termed it as C19Q-KRAB. C19Q-KRAB could presumably recruit SETDB1 [24, 25] which deposits loci-specific H3K9me3 [26]. HP1α could then interact with loci or regions with H3K9me3 [27, 28]. Thus, we expect that EpiGo of C19Q will allow us to track the changes of genome organization upon epigenetic alterations in living cells. To further track the dynamic interaction between EpiGo-mediated H3K9me3 and HP1α condensates, HaloTag was knocked-in at the C-terminus of HP1α by CRISPR-Cas9 system. This U2OS-HP1α-HaloTag cell line was then used to generate cell lines stably expressed C19Q-sgRNA-2XPP7, PCP-GFP and dCas9 or dCas9-KRAB, resulting in U2OS-Epi Go-Control or U2OS-EpiGo-KRAB cells for direct visualization of C19Q upon ectopic H3K9me3 modifications (**Figure 1B and 1C**). Finally, ChIP-seq analysis confirmed that most target site of C19Q successfully acquired ectopic H3K9me3 (**Fig. S1 and Fig. 1D**). These data demonstrate that we have successfully induced large scale epigenetic alterations using EpiGo-KRAB.

**Fig. 1.**
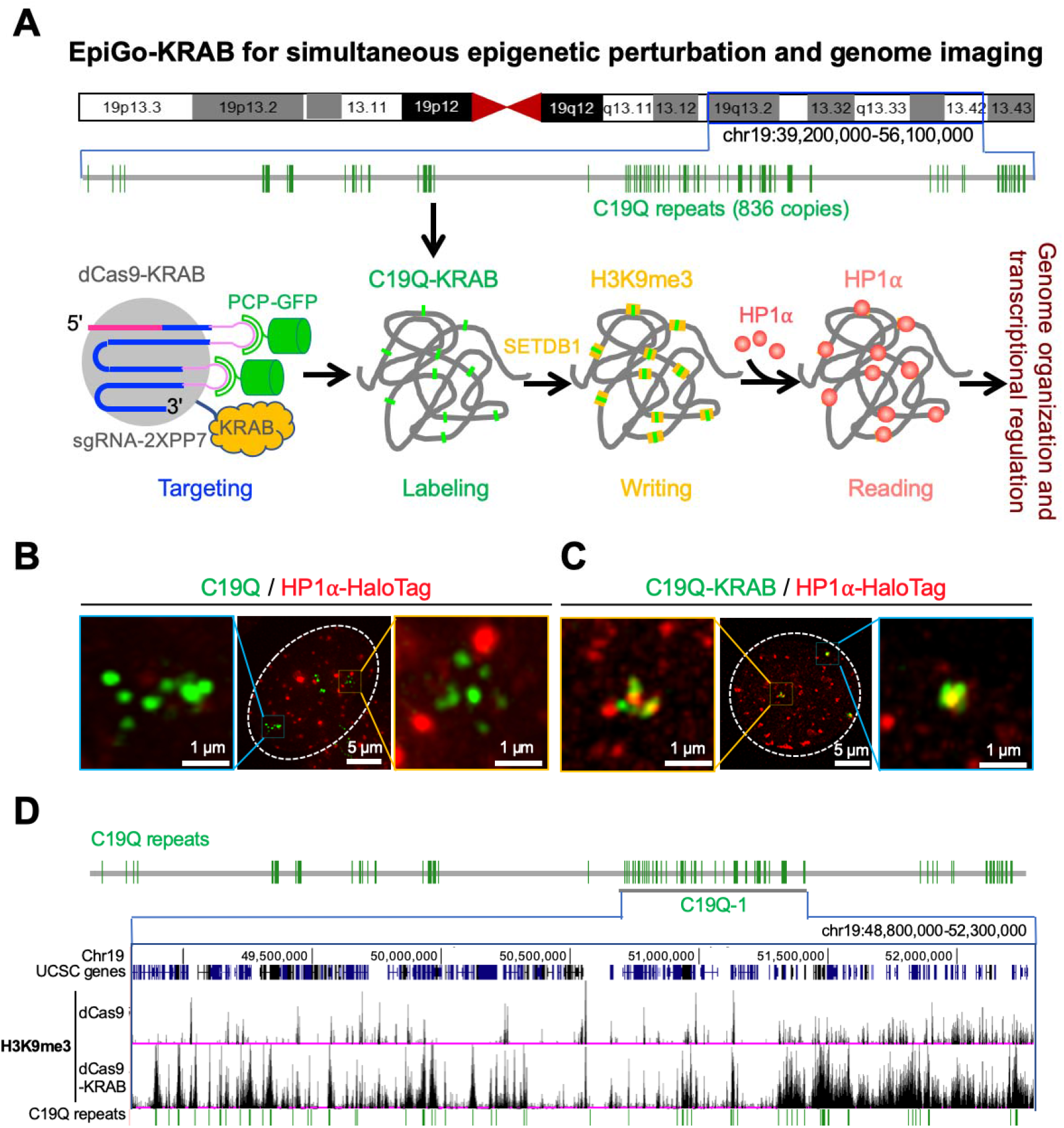
A CRISPR-based EpiGo-KRAB system to track genome organization upon epigenetic modification. (A) A scheme of the EpiGo system. EpiGo (<u>E</u>pigenetic <u>p</u>erturbation <u>i</u>nduced <u>G</u>enome <u>o</u>rganization) system consists of dCas9-KRAB, PCP-GFP and sgRNA-2XPP7 targeting to C19Q repeats, which contain 836 copies of target sites on the q-arm of chromosome 19. C19Q-KRAB will recruit SETDB1 and induce H3K9 trimethylation at each target sites. H3K9me3 regions will recruit HP1α and induce genome organization. (B) CRISPR-based imaging of C19Q in presence of PCP-GFP, C19Q-sgRNA-2XPP7 and dCas9, marked with GFP-C19Q. The HP1α-HaloTag stable cell lines was used to examine the colocalization between C19Q (green) and HP1α (red). (C) CRISPR imaging with the same conditions as (B) except that dCas9 was replaced by dCas9-KRAB and marked with GFP-C19Q-KRAB. (D) H3K9me3 state of EpiGo-Control (dCas9) and EpiGo-KRAB (dCas9-KRAB) cell lines. ChIP-seq of H3K9me3 was performed in these two cell lines. C19Q-1 region (chr19:48,800,000-52,300,000) is chosen to show the difference of H3K9me3 state between EpiGo-Control and EpiGo-KRAB cell lines.

### EpiGo-KRAB induces SETDB1-dependent genomic clustering and phase separation

We then sought to examine the spatiotemporal dynamics of C19Q regions upon EpiGo-KRAB mediated H3K9me3. As shown in **Fig. 2A**, discrete foci of C19Q (green) were visible by CRISPR labeling in U2OS-EpiGo-Control (dCas9) cells, which barely colocalize with HP1α (red) over time. However, In EpiGo-KRAB cell lines (dCas9-KRAB), C19Q loci dynamically interacted with HP1α condensates and clustered at the surface of HP1α condensates. Eventually, adjacent HP1α condensates coalesced together. To investigate how HP1α mediates local compaction of genomic regions upon EpiGo-KRAB induction (**Fig. 2B**), structured illumination microscopy (3D-SIM) was used to acquire high resolution imaging of C19Q and HP1α. We found that C19Q foci decorated on the surface of HP1α condensates in EpiGo-KRAB (dCas9-KRAB) cell lines (**Fig. 2B**). Quantitative analysis confirmed that C19Q foci were not clustered in EpiGo-Control cells, while 90% of C19Q foci showed clustering in EpiGo-KRAB cells (**Fig. 2C**). In sum, these results suggest that EpiGo-KRAB can induce genome reorganization through a dynamic and stepwise process.

**Fig. 2.**
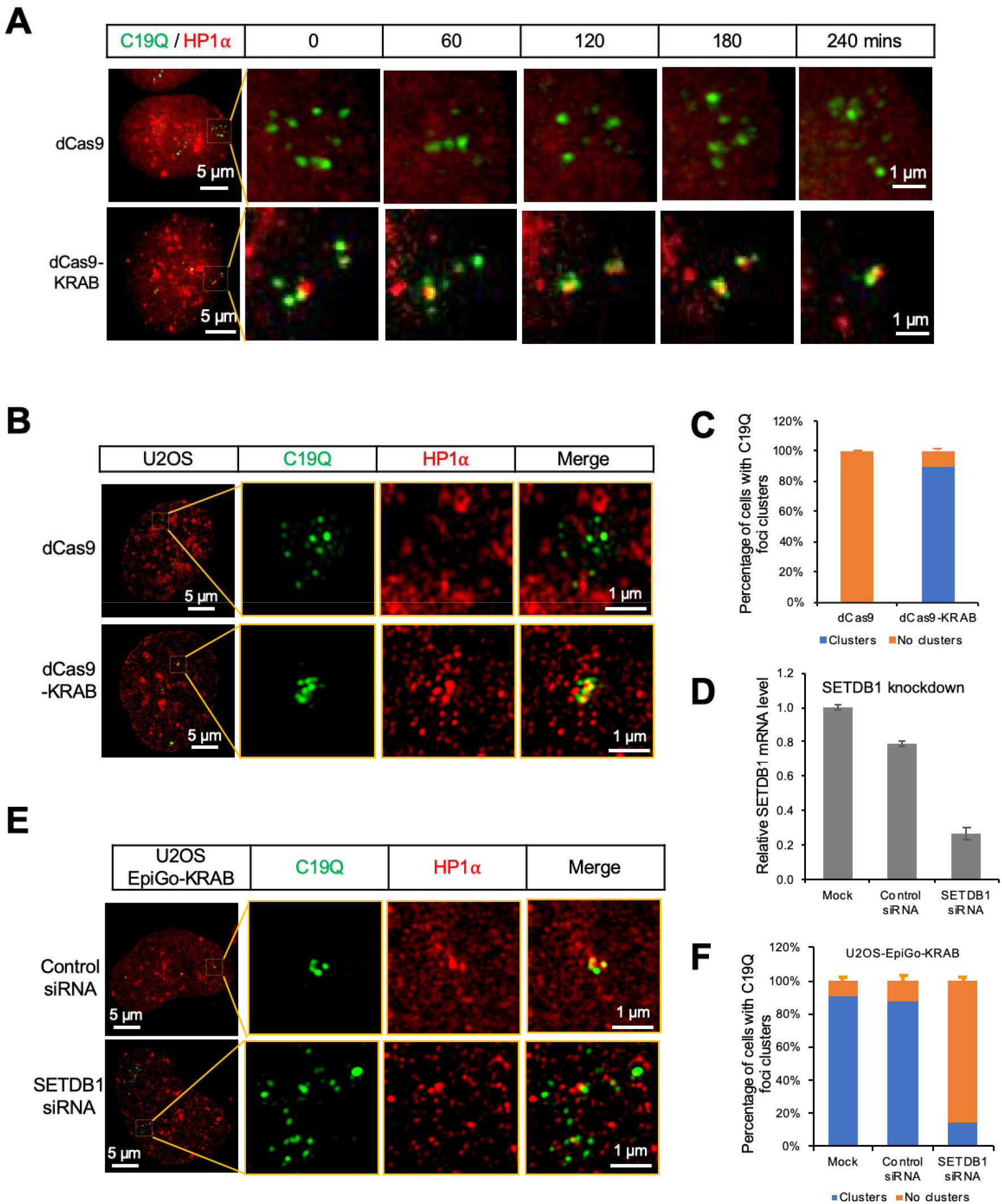
SETDB1 is required for EpiGo-KRAB mediated genomic clustering. (A) Live cell tracking of C19Q dynamics in EpiGo-Control (dCas9) and EpiGo-KRAB (dCas9-KRAB) cell lines. These two cell lines were stably expressed PCP-GFP, C19Q-sgRNA-2XPP7 and dCas9 or dCas9-KRAB. The HaloTag was knocked-in at the C-terminus of HP1α and used to examine the colocalization between C19Q (green) and HP1α (red). The dCas9 or dCas9-KRAB was induced to express 24 hours before tracking. The dynamics of C19Q (green) and HP1α (red) were tracked for 240 mins. Images were captured every 30 mins. (B) 3D-SIM images of C19Q and HP1α in EpiGo-Control and EpiGo-KRAB cell lines. C19Q (green) was labelled by GFP and HP1α (red) was labelled by HaloTag. The colocalization between C19Q and HP1α was shown in merged images. (C) Percentage of C19Q foci clusters in EpiGo-Control (dCas9) and EpiGo-KRAB (dCas9-KRAB) cell lines. Data are presented as means ± SD (n=3). (D) RT-qPCR analysis of SETDB1 knockdown efficiency. RNA was extracted from the EpiGo-KRAB cells lines transfected with either control siRNA or SETDB1 siRNA. Data are presented as means ± SD (n=3). (E) 3D-SIM images of C19Q (green) and HP1α (red) in EpiGo-KRAB cell lines in the presence of control siRNA or SETDB1 siRNA. (F) Percentage of C19Q foci clusters in the U2OS-dCas9-KRAB cells lines transfected with either control siRNA or SETDB1 siRNA. Data are presented as means ± SD (n=3).

Immunofluorescence also showed that H3K9me3 surrounded the HP1α condensates (**Fig. S2**). It was reported that dCas9-KRAB mediated H3K9me3 deposition via histone methyltransferase SETDB1 [25, 29]. To test whether SETDB1 is essential for EpiGo-KRAB mediated genomic clustering, SETDB1 was knocked down by siRNAs in EpiGo-KRAB (dCas9-KRAB) cell lines (**Fig. 2D)**. The mRNA levels of SETDB1 decrease to 26% of the mock cells when transfected with SETDB1 siRNA. The percentage of C19Q clustering is 87% in cells transfected with control siRNA, but markedly decreased to 14% in cells transfected with SETDB1 siRNA (**Fig. 2F**). Finally, C19Q loci were highly dynamic in EpiGo-Control cells but the mobility of C19Q loci dramatically decreased in EpiGo-KRAB cell lines (**Fig. S3**). These results support a role of H3K9me3 in mediating phase separation of HP1α droplets and large-scale genome organization.

### EpiGo-KRAB mediated H3K9me3 and genomic clustering does not result in widespread gene silencing

We then sought to investigate the relationship of ectopic H3K9me3 and transcription. The C19Q repeats are present in gene body, promoter or distal regions (**Fig. S4A**). The C19Q-1 region spans 3.5 Mb and contains 146 genes (**Fig. 3A**). This region contains clearly two sub-regions: regions 1 and 2 (**Fig. 3A and Fig. S4B–4C**). Most genes (64%) in Region 1 are active or modestly active, while the majority of genes (79%) in Region 2 are inactive in EpiGo-Control (dCas9) cells. As shown in **Fig. S4D**, EpiGo-KRAB does not affect the global expression levels. A close examination of EpiGo-KRAB targeted genes revealed that a small set of active genes are indeed silenced. For these genes, such as the top five silenced genes *(KCNN4, KDELR1, HSD17B14, KLK6* and *TNNT1),* ectopic H3K9me3 primarily occurred at the promoter regions, which is accompanied by the loss of H3K4me3 and H3K27ac (**Fig. 3B-3C and Fig. S4E**). This observation was confirmed by global analysis of all genes targeted by EpiGo-KRAB (**Fig. 3D–3F**). Unexpectedly, we found about the expression of the majority (61 out of 73) of active genes in Region 1 and 2 was not affected by EpiGo-KRAB (**Fig. 3A and Fig. S4B–4C**). For example, *CYTH2,* which acquired ectopic H3K9me3 in the 3’-UTR region, did not show significant changes of promoter chromatin states and transcriptional level (**Fig. 3B, 3D–3F**). These results suggest that EpiGo-KRAB mediated global chromatin clustering does not result in widespread gene silencing beyond its direct promoter targets.

**Fig. 3.**
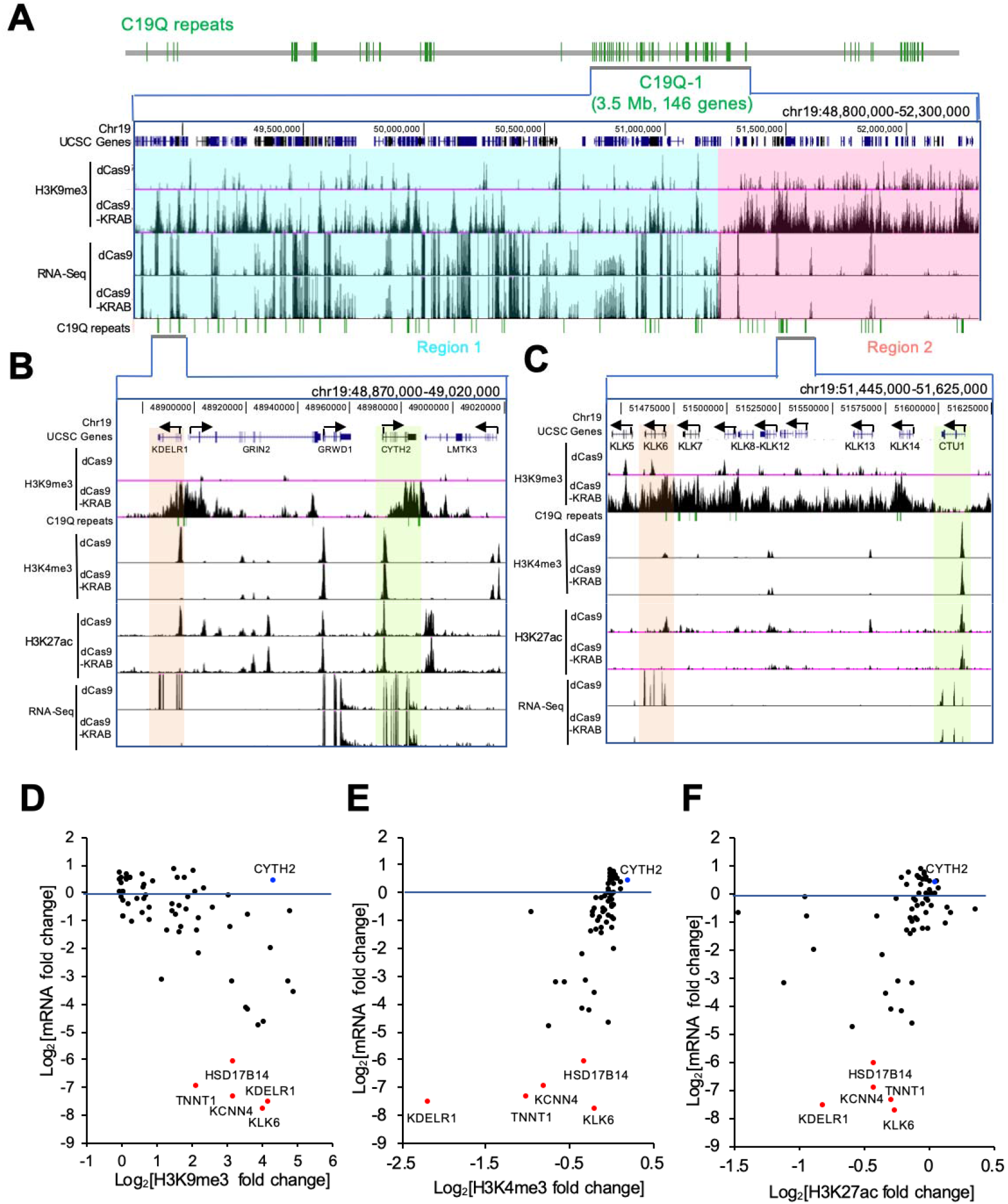
EpiGo-KRAB mediated H3K9me3 deposit does not result in widespread gene silencing. (A) H3K9me3 and RNA levels in EpiGo-Control (dCas9) and EpiGo-KRAB (dCas9-KRAB) cell lines. ChIP-seq of H3K9me3 and RNA-seq were performed in these two cell lines. Region 1 (chr19: 48,800,000-51,200,000, cyan) or Region 2 (chr19:51,200,001-52,300,000, pink) of C19Q-1 with different transcriptional states was highlighted with cyan or pink respectively. (B) H3K9me3, H3K4me3, H3K27ac states and RNA level in Region 1. ChIP-seq of H3K9me3, H3K4me3, H3K27ac and RNA-seq were performed in EpiGo-Control (dCas9) and EpiGo-KRAB (dCas9-KRAB) cell lines. *KDELR1* gene or *CYTH2* gene were highlighted with light orange or light green respectively. (C) H3K9me3, H3K4me3, H3K27ac states and RNA levelin Region 2. *KLK6* gene or *CTU1* gene were highlighted with light orange or light green respectively. (D) Fold changes of mRNA level and H3K9me3 in the EpiGo-KRAB targeted genes. The active genes (FPKM ≥1) targeted by EpiGo-KRAB were chosen for the analysis. The five genes *(KCNN4, KDELR1, HSD17B14, KLK6* and *TNNT1)* with most reduction in mRNA levels were marked in red. (E) Fold changes of mRNA level and H3K4me3 in the EpiGo-KRAB targeted genes. (F) Fold changes of mRNA level and H3K27ac in the EpiGo-KRAB targeted genes.

Notably, the peaks of EpiGo-induced ectopic H3K9me3 were negatively correlated with active transcription **(Fig. 3B)**, as also observed for endogenous H3K9me3 genome wide (**Fig. S5**). Interestingly, ectopic H3K9me3 appears to frequently exist as continuous domains in Region 2 (**Fig. 3C**). For example, H3K9me3 spreads beyond C19Q repeats for hundreds of kilobases in Region 2 until the proximity of an active gene *CTU1* (**Fig. 3C**). By contrast, ectopic H3K9me3 in active region (Region 1 in **Fig. 3B**) is intermittent and its spreading is frequently restricted, often stopping at the sites of active genes. These results raise a possibility that H3K9me3 spreads more efficiently in transcriptionally silenced regions but may be antagonized by active chromatin states.

### EpiGo-KRAB induces *de novo* heterochromatin-like domain formation

To further examine genome organization of the active or silenced regions upon H3K9me3, Oligopaint FISH was used to visualize Region 1, Region 2 (**Fig. 4A and Table S2-S3**). Region 1 was transcriptionally active and showed intermittent H3K9me3 in EpiGo-KRAB cell lines. Region 2 was silenced and showed more continuous H3K9me3 in EpiGo-KRAB cell lines. We also included a neighbor region 3’ downstream of Region 2 as a comparison (Region 3). Region 3 showed strong and continuous H3K9me3 in both EpiGo-Control and EpiGo-KRAB cell lines, consistent with the fact that it is not targeted by EpiGo-KRAB. In EpiGo-Control cells, Region 1 showed multiple discrete foci, which partially overlap with C19Q probe but barely associate with HP1α (**Fig. 4B)**. By contrast, Region 2 exists as only one focal spot in the majority of cells (**Fig. 4B–4C**). It overlaps with C19Q but, surprisingly, barely associated with HP1α, suggesting that Region 2 is compacted independent of HP1α association before EpiGo-KRAB induction and ectopic H3K9me3 acquisition. Region 3, which is extensively modified by H3K9me3 in EpiGo-KRAB cells, has one focal spot and is associated with HP1α (**Fig. 4B–4C**). These results suggested that silenced regions (Region 2 or 3) are more compacted than active regions (Region 1), and their association with HP1α appears to be related to H3K9me3 strength. Indeed, in EpiGo-KRAB cells, Region 1 collapsed into one focal spot and associated with HP1α (**Fig. 4D–4E**). Region 2 again had only one focal spot, but now became associated with HP1α. These results indicated that EpiGo-KRAB promotes its association with HP1α condensates and further chromatin compaction perhaps in an H3K9me3-density dependent manner.

**Fig. 4.**
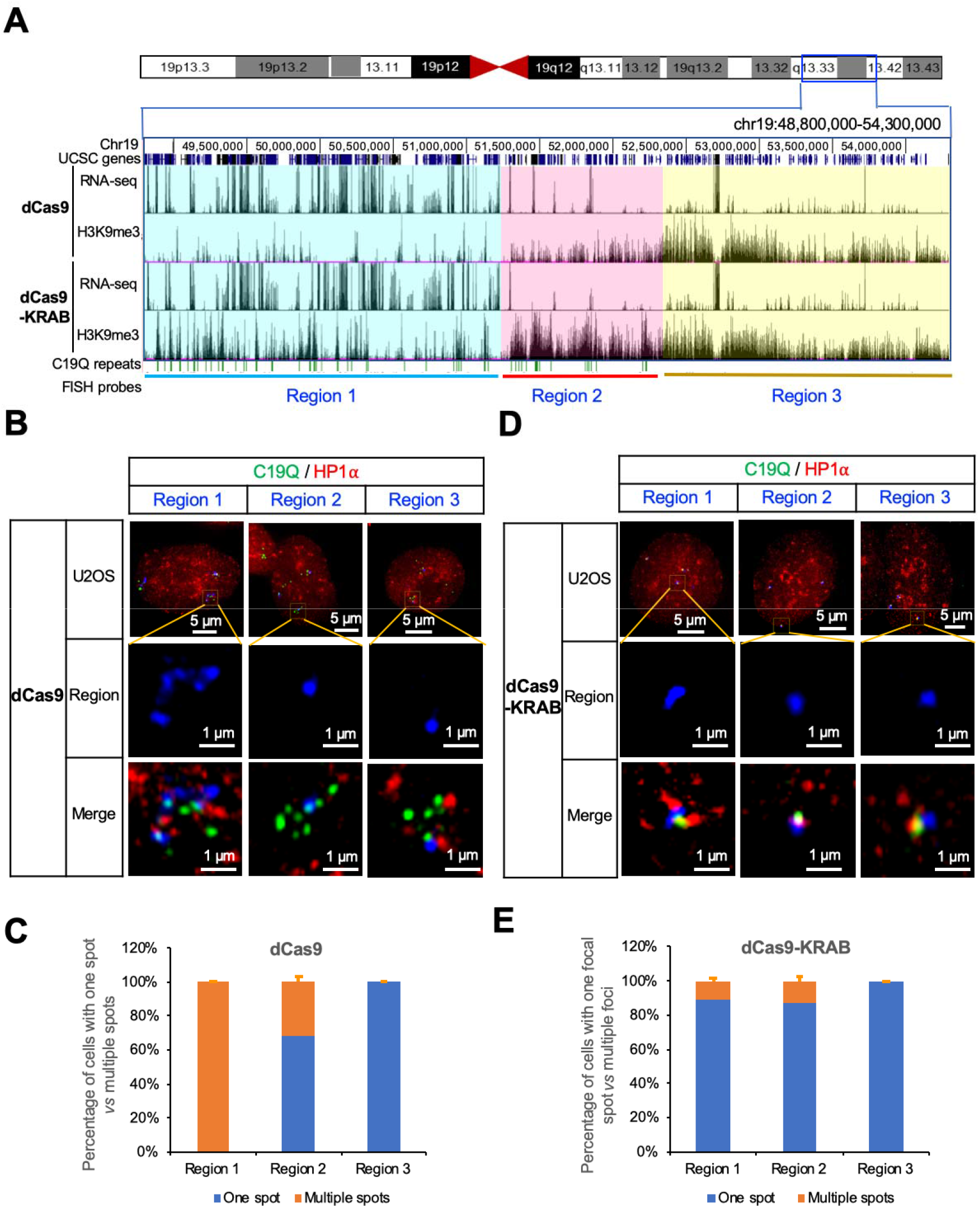
EpiGo-KRAB induced *de novo* heterochromatin-like domain formation. (A) H3K9me3 states and RNA Levels of Region 1, Region 2 and Region 3 in EpiGo-Control (dCas9) and EpiGo-KRAB (dCas9-KRAB) cell lines. Region 1 (chr19: 48,800,000-51,200,000) representing an active region, Region 2 (chr19: 51,200,001-52,300,000) representing a silenced region, Region 3 (chr19: 52,300,001-54,300,000) containing endogenous H3K9me3 was highlighted in cyan, pink and yellow respectively. The target sites of C19Q repeats were shown in green. Oligopaint FISH probes were designed to detect Region 1, Region 2 and Region 3 respectively. (B) Images of C19Q, HP1α and Region 1, 2 or 3 in EpiGo-Control cell lines. C19Q (green) was labelled by GFP using CRISPR imaging, HP1α (red) was labeled by HaloTag and Region 1, 2 or 3 (blue) was labelled by in situ hybridization with DNA probes conjugated to the dye Alexa-647. The colocalization among C19Q, HP1α and Region 1, 2 or 3 was shown in merged images. (C) Percentage of cells with one spot vs multiple spots in Region 1, 2, or 3 of EpiGo-Control (dCas9) cell lines. Data are presented as means ± SD (n=3). (D) Images of C19Q, HP1α and Region 1, 2 or 3 in EpiGo-KRAB cell lines. All the conditions are the same as (B) except the dCas9 was replaced by dCas9-KRAB. (E) Percentage of cells with one spot vs multiple spots in Region 1, 2, or 3 of EpiGo-KRAB (dCas9-KRAB) cell lines. Data are presented as means ± SD (n=3).

### EpiGo-KRAB induced large-scale rearrangement of chromatin compartmentalization

Genome is organized in 3D from sub-kilobase to megabase scales and segregated into domains in cellular nucleus [4]. Genome compartmentalization occurs at the hundreds of kilobases to megabase scales from Hi-C heatmap [11]. To understand how epigenetic manipulation affects compartmentalization in active and silenced regions, Hi-C was performed on U2OS-EpiGo-Control (dCas9) and U2OS-EpiGo-KRAB (dCas9-KRAB) cell lines. As a control, no drastic changes of global compartmentalization were observed in the Hi-C matrix of the genome upon EpiGo-KRAB induction (data not shown). Focused on the C19Q region, which showed substantial changes of H3K9me3 states, we found drastic rearrangement of local compartments (**Fig. 5A and 5B**). For instance, the entire Region 2 and Region 3 fused to become a large compartment in EpiGo-KRAB cells, which even erodes part of Region 1 (1/2/3 compartment) (**Fig. 5B**). Throughout the regions, we have found a number of regions with C19Q repeats that show increased interactions with their neighbor regions, leading to compartment merging (red arrows, **Fig. 5A**). These results indicated that EpiGo-KRAB induced extensive rearrangement of chromatin compartments. We reasoned that this is possibly because H3K9me3 marked regions anchor chromatin to HP1α condensates, which breaks existing compartmentalization [30] and form new compartments. Therefore, the final compartments in the genome may be decided by the overall topology executed through a default compartmentalization (gene density and transcriptional state-correlated) altered by architectural proteins and epigenetic states.

**Fig. 5.**
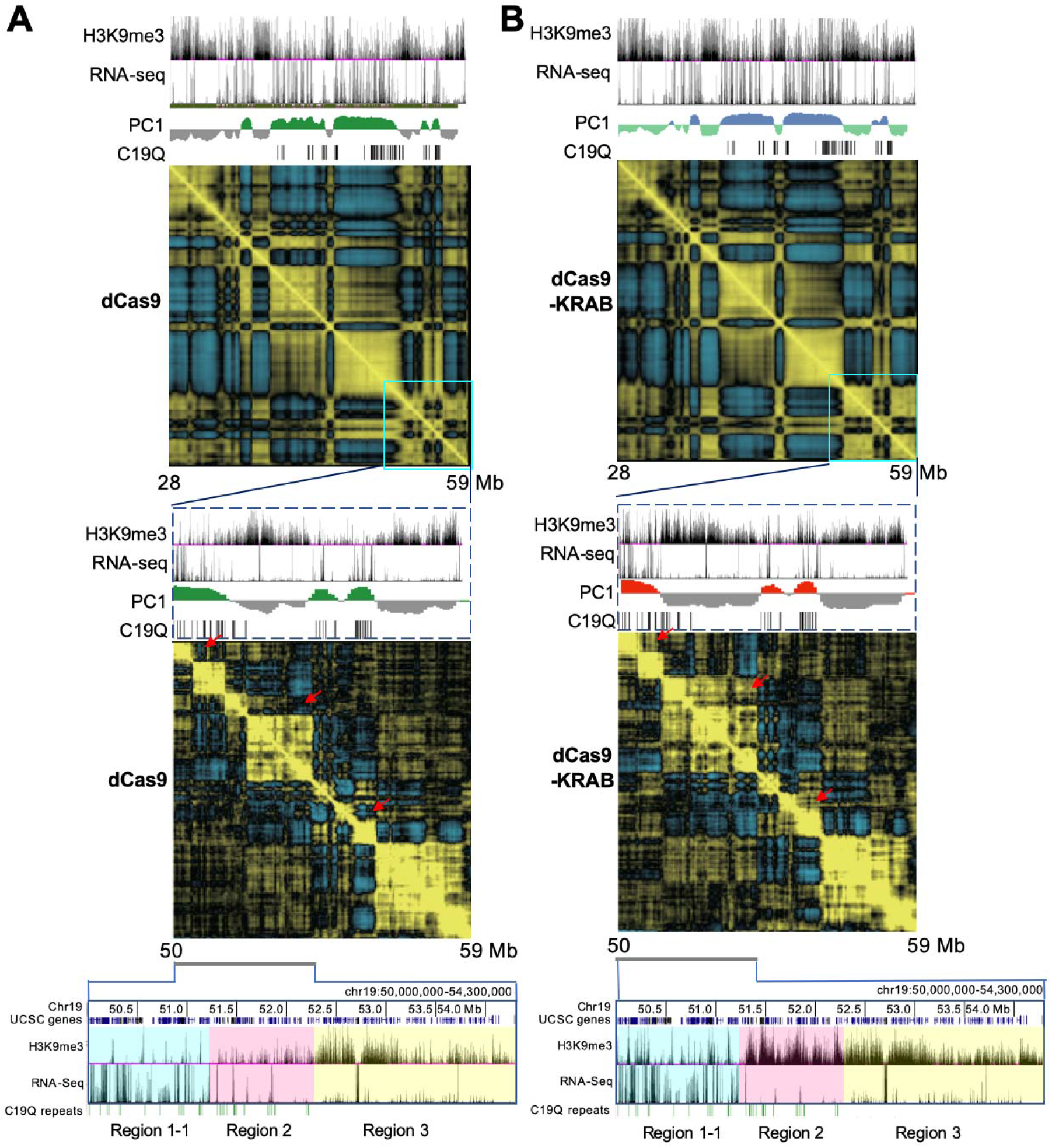
EpiGo-KRAB induced rearrangement of chromatin compartmentalization. (A) Hi-C heatmap shown the compartmentalization of the C19Q region in EpiGo-Control (dCas9) cells. H3K9me3 state, RNA level, PC1 and C19Q repeats (target sites) shown at the top of the Hi-C heatmap (Chr19: 28,000,000-59,000,000). Hi-C heatmap of a local region (chr19: 50,000,000-59,000,000) was used to show the local compartmentalization. Region 1-1 (a partial Region 1), Region 2 and Region 3 were shown in cyan, pink and yellow respectively. (B) Hi-C heatmap shown the compartmentalization of the C19Q region in EpiGo-KRAB (dCas9) cells. All the data processing and display are the same as (A).

## Discussion

Despite the long-observed correlation, whether epigenetic modifications can regulate 3D genome architecture in live cells remains poorly understood. The EpiGo-KRAB system provides a powerful tool to allow H3K9me3 at specific genes or regions and track their changes of location, structure and dynamics. Our data show that EpiGo-KRAB is able to mediate *de novo* heterochromatin-like domain formation in a dynamic and stepwise process (**Fig. 6**): 1) EpiGo-KRAB induces ectopic H3K9me3 at defined loci which further spread depending on the existing chromatin states and transcriptional activities; 2) EpiGo-KRAB mediated H3K9me3 promotes chromatin anchoring to HP1α condensates; 3) EpiGo-KRAB mediated heterochromatin-like domain, possibly by phase separation. These results also suggest neither non-promoter ectopic H3K9me3 *per se* nor H3K9me3-mediated chromatin compaction and phase separation with HP1α condensates is sufficient for gene silencing. In fact, active transcription can persist in EpiGo-KRAB induced cells and appear to be able to antagonize H3K9me3 spreading. It would be worthy to further explore how transcription and H3K9me3 interplay in space and time with diverse biological context.

**Fig. 6.**
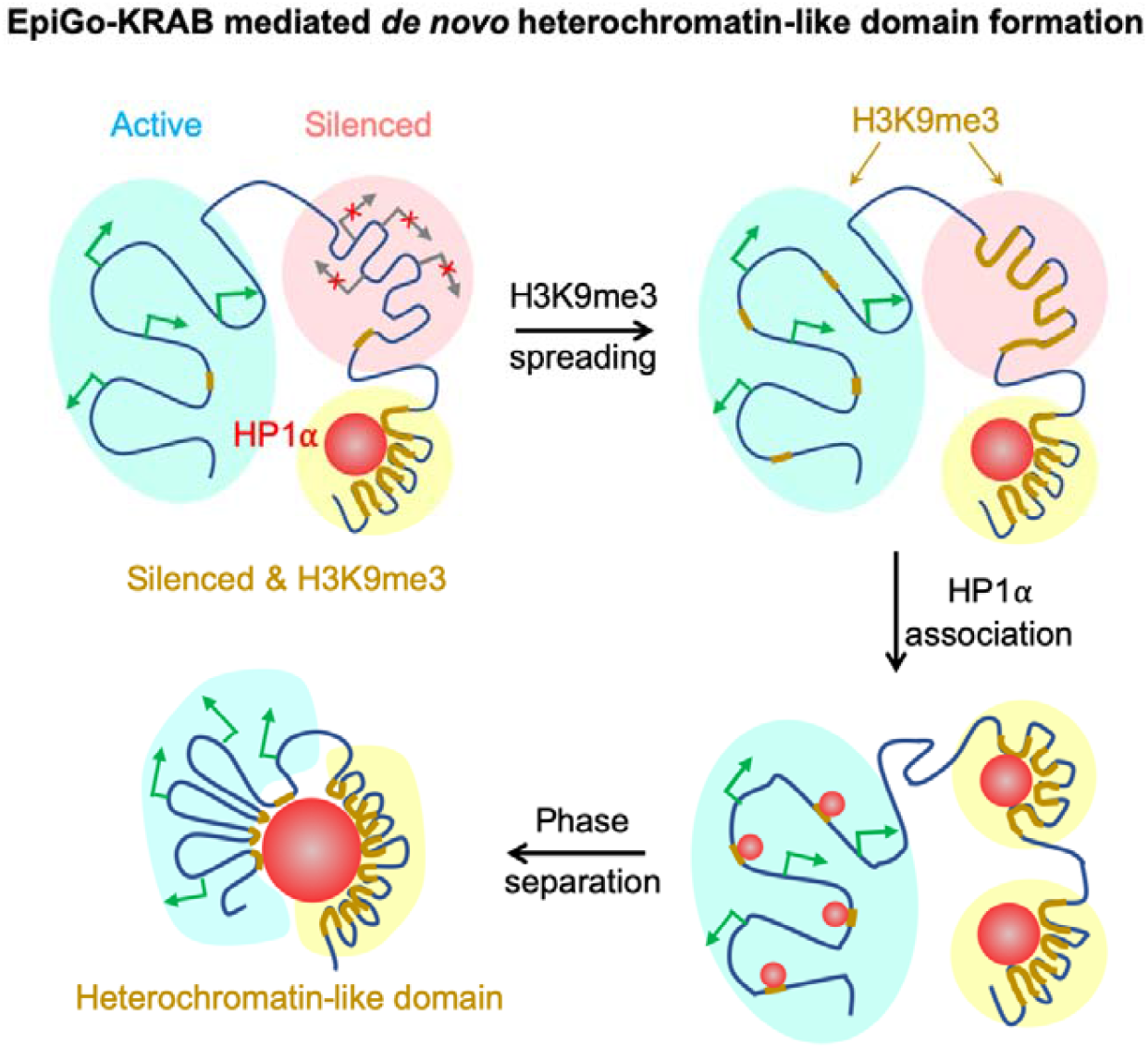
A model for genome organization mediated by H3K9me3 in living cells. H3K9me3 spread continuously over hundreds of kilobases to megabases in silenced regions, but the spreading frequently stops at the transcriptional initiation sites in active regions. Loci or regions with H3K9me3 decorate on the surface of HP1α condensates, which mediates chromatin local compaction. H3K9me3 and local compaction does not result in widespread gene silencing in active regions, but induces large scale chromatin compartmentalization possibly by phase separation.

Another interesting question is what degree of H3K9me3 is required for HP1α condensate association, chromatin local compaction or genome compartmentalization. Here, by analyzing different regions with distinct transcriptional activities, we found that repressed regions can form compact DNA without associating with HP1α condensate, which is consistent to recent finding that compaction of mouse heterochromatin foci can be independent of HP1 [31]. However, high levels of ectopic H3K9me3 may be necessary to promote HP1α association and chromatin compaction in active regions.

Finally, drastic changes of epigenetic modifications such as H3K9me3, H3K27me3 and DNA methylation occur in stem cell differentiation, embryonic development and many diseases [32–34]. With the EpiGo system, it will be interesting to explore how tissue-specific epigenetic modifications are established and maintained at kilobase to megabase scales, and how they regulate chromatin architecture, gene expression and cell fate decision.

## Methods

### Cell culture

The U2OS (human bone osteosarcoma epithelial, female) cell, and HEK293T cells (human embryonic kidney epithelial, female) were cultured in DMEM (Life Technologies) with high glucose in 10% FBS (fetal bovine serum, Life Technologies). All cells were cultured at 37°C and 5% CO_2_ in a humidified incubator.

### Chromosome-specific repeats for the EpiGo system

Mining of chromosome-specific repeats was described previously with some modifications [35]. The human reference genome (assembly GRC h37/hg19) was downloaded from the UCSC Genome Browser (http://genome.ucsc.edu) to find target regions and design guide RNAs. The bioinformatics tool Jellyfish [36] was used to find all 15-mers (12-mers ending with NGG or starting with CCN) with at least 5 copies in any chromosome. The 15-mers with more than 100,000 targets were filtered out. Each candidate 15-mer was searched for off-targets in all other chromosomes. The candidate 15-mer was discarded if there was any cluster of 5 off-targets or more within any 50 kb region. The C19Q repeats (**Table S1**), which consists of 836 copies of CRISPR target sites spanning ~17 megabases at the q-arm of chromosome 19, was chosen as EpiGo targets in this study. The overlapping genes including the intron/exon information were extracted from the GENCODE Genes track (version 27lift37), downloaded from the UCSC Genome Browser.

### Plasmids construction

The expression plasmid pHAGE-TO-dCas9 has been described previously [19, 23], in which HSA-P2A was inserted at the N-terminal of dCas9 resulting in pHAGE-TO-HSA-P2A-dCas9 and KRAB was then subcloned into the C-terminal of dCas9 resulting pHAGE-TO-HSA-P2A-dCas9-KRAB. The expression plasmid pHAGE-EFS-PCP-GFP has been described previously [23]. The expression vector for guide RNA was based on the pLKO. 1 lentiviral expression system, in which TetR-BFP-P2A-2XPP7 was inserted right after the phosphoglycerate kinase (PGK) promoter, resulting pTetR-P2A-BFP-2XPP7. The expression plasmid for guide RNAs targeting to C19Q was made by using the rapid guide RNA construction protocol described previously [35].

The donor plasmid for knock-in of HaloTag at the C-terminal of HP1α was made by Golden Gate cloning. The donor consists of three fragments: A 400-bp fragment just upstream of stop codon of HP1α (left arm), HaloTag coding sequences and a 400-bp fragment just downstream of stop codon of HP1α (right arm). Left arm and right arm fragments were amplified from U2OS genomic DNA isolated with Cell DNA isolation mini kit (Vazyme). The HP1α-HaloTag donor was assembled by Golden Gate cloning method into pDONOR vector [35], resulting in pDONOR-HP1α-HaloTag. The guide RNA targeting sequences AAACAGCAAAGAGC<u>TAA</u>AGG spanning the stop codon (TAA, underlined) of HP1α and cloned into guide RNA expression vector pLH-sgRNA2 [23], resulting in pLH-sgRNA2-HP1α.

### Generation of U2OS-HP1α-HaloTag cell lines

For knock-in of HaloTag into HP1α locus, U2OS cells were co-transfected with 200 ng of pHAGE-TO-Cas9, 600 ng of pLH-sgRNA1-HP1α and 600 ng pDONOR-HP1α-HaloTag using Lipofectamine 2000 (Life Technologies) for 6 hours and then replaced culture media containing 2 nM HaloTag-JF549. The transfected cells were cultured for additional 24-48 hours before examining the knock-in efficiency by fluorescent imaging or flow cytometry.

Fluorescence imaging was used to check the proper localization of HP1α-HaloTag. The HaloTag positive cells was sorted by BD FACS Aria III equipped with 561 nm excitation laser, and the emission signals were detected by using filter at 610/20 nm (wavelength/bandwidth) for HaloTag-JF549. The localization of HP1α-HaloTag was examined again under fluorescence microscope after cultured for additional two weeks. The resulting cell line was named U2OS-HP1α-HaloTag.

### Generation of U2OS-EpiGo cell lines

For generation of U2OS-EpiGo cell lines, lentivirus for PCP-GFP, sgRNA, dCas9, or dCas9-KRAB was used. HEK293T cells were seeded into 6-well and transfected with 0.5 μg pCMV-dR8.2-dvpr, 0.3 μg pCMV-VSV-G and 1.5 μg of transfer plasmid (pHAGE-EFS-PCP-GFP, pTetR-P2A-BFP-sgRNA1-2XPP7-C19Q, pHAGE-TO-dCas9 or pHAGE-TO-dCas9-KRAB) using Lipofectamine 2000 (Invitrogen) following manufacturer’s protocol. After 48 hours, supernatant was harvested and filtered with 0.45-μm polyvinylidene fluoride Syringe filters. The filtered supernatant was titrated, and either directly used or concentrated. The concentration was performed by using Lentivirus Concentration Reagent (Biodragon-immunotech. Inc). The concentrated viral particles were immediately used for transduction or stored at −80°C in aliquots.

U2OS-HP1α-HaloTag cells were grown to 40% confluency in 6-well plates and viral supernatants for PCP-GFP, sgRNA targeting to C19Q, dCas9 or dCas9-KRAB were added for transduction. Fresh media containing 500 ng/ml doxycyline (Sigma-Aldrich) was used 48 hours post-infection. After additional 24 hours, Alexa Fluor^®^ 647 anti-mouse CD24 Antibody (BioLegend) for HSA were added before cell sorting. Expression levels of PCP-GFP, sgRNA (BFP) and dCas9 (AlexaFluor 647) or dCas9-KRAB (AlexaFluor 647) was examined by BD FACS Aria III equipped 405 nm, 488 nm, and 647 nm excitation lasers, and the emission signals were detected by using filters at 450/40 nm (wavelength/bandwidth) for BFP, 530/30 nm for GFP and 662/20 nm for AlexaFluor 647. To generate single colonies, single cells with different levels of PCP-GFP, sgRNA and dCas9 or dCas9-KRAB were sorted into 96-well plates. After 10-14 days, triple positive colonies were further examined for CRISPR-based labeling of C19Q foci under fluorescence microscope. The C19Q labeled cells were propagated and named U2OS-EpiGo-Control (PCP-GFP, C19Q sgRNA and dCas9) and U2OS-EpiGo-KRAB (PCP-GFP, C19Q sgRNA and dCas9-KRAB) respectively.

### Oligopaint FISH probe design and preparation

Oligopaint FISH probes targeting the q-arm of chromosome 19 were designed according to OligoMiner (https://github.com/brianbeliveau/OligoMiner) and probe datasets for Human chromosome 19 from hg19 with mining settings of Balance (https://oligopaints.hms.harvard.edu/genome-files). Here we focused on the region of chr19: 48.8-54.3 Mb (hg19) including Region A (chr19: 48.8-51.2 Mb), Region B (chr19: 51.2-52.3 Mb) and Region C (chr19: 52.3-54.3 Mb). The detail design of Region A, Region B and Region C probes were shown in **Table S2** and **Table S3**. The oligo pool for Oligopaint FISH probes was ordered from Synbio Technologies Inc. The preparation of Oligopaint FISH probes has been described previously [14, 37]. Briefly, the oligo pools for Region A, Region B and Region C were amplified via limited-cycle PCR (Vazyme), with forward primer and corresponding reverse readouts (one per 300 kb region) with T7 promoter sequence TAATACGACTCACTATAGGG appended to its 5’ end. Each pool was purified by AxyPrep DNA Gel Extraction Kit (Axygen). The oligo pools for Region A, Region B or Region C were used as templates for in vitro transcription by HiScribe T7 quick high yield RNA synthesis kit (NEB). The obtained RNA products were converted back to DNA oligo probes via reverse transcription (RT) with common RT primer (ATCGACCCGGCATCAACGCCACGATCAGCT) conjugated with a 5’ AlexaFluor 647 using Maxima H Minus RT enzyme (Thermo Fisher). The intermediate RNA products were removed with NaOH-EDTA solution in 95 °C for 10 minutes and the single strand DNA oligo probes were purified via DNA clean & concentrator-25 (D4033, Zymo Research). The ssDNA probes were either directly used for Oligopaint FISH or store in aliquots at −80°C.

### Immunofluorescence

Cells were cultured on #1 coverslip (Electron Microscopy Sciences), and then fixed with 4% PFA for 10 min, washed three times with 1XPBS and permeabilized by 0.1% Triton X-100 in 1XPBS for 10 min. Cells were incubated with rabbit polyclonal antibodies against H3K9me3 (1:1000; Active Motif) in the 1XPBS for 1 hour at room temperature. After washed three times with 1XPBS and incubated with Goat anti-rabbit lgG (H+L) secondary antibody (1:500, Invitrogen) for 1 hours. After washed three times with 1XPBS, coverslips were mounted with Prolong antifade reagents (Invitrogen) for imaging.

### ImmunoFISH

1×10^5^ of U2OS-EpiGo cells were cultured on the #1 coverslip (Electron Microscopy Sciences) and added 500 ng/ml doxycycline for induction of dCas9 or dCas9-KRAB expression. After 72 hours, cells were fixed with 4% PFA for 10 minutes, washed by 1XPBS for three times and permeabilized with PBS-0.5%Triton X-100 for 10 minutes.

Cells were incubated with Rabbit anti-GFP polyclonal antibody (ab290, Abcam) in the 1XPBS for 1 hour at room temperature. After washed three times with 1XPBS and incubated with Alexa Fluor 488 Goat anti-rabbit lgG (H+L) secondary antibody (1:500, Invitrogen) for 1 hour. After washed three times with 1XPBS, cells were fixed with 4% PFA again for 20 min. The fixed cells were treated with 0.1N HCl for 5 min and washed with 2XSSCT (300 mM NaCl, 30 mM sodium citrate and 0.1% Tween-20), followed by incubation with 50% (vol/vol) formamide in 2XSSCT in 60°C for 20 min. Samples were denatured at 78°C for 3 min with 20 μl hybridization buffer (2XSSC, 50% formamide, 10% dextran sulfate, 0.2% Tween-20 mixed with ~1.6 μM total DNA oligo probes) and incubated at 42°C for 16 hours in a humidified chamber. Samples were washed by 2XSSCT for 4 times at 60°C and 2 times at room temperature. The cells were washed by 1XPBS with 0.5 M NaCl before mounted with Prolong antifade reagents (Invitrogen) for imaging.

### Live cell imaging

All live cell imaging was carried out on a DeltaVision OMX^TM^ V4 imaging system (GE Healthcare), equipped with a 60x (NA 1.42) Plan Apo oil-immersion objective (Olympus). The cells were cultured on No. 1.0 glass bottom dishes (MatTek). The microscope stage incubation chamber was maintained at 37°C and 5% CO_2_. HaloTag-JF549 was excited at 561 nm, and its emission was collected using filter at 609/37 nm (wavelength/bandwidth), GFP was excited at 488 nm and collected using filter at 498/30 nm (wavelength/bandwidth). Imaging data were acquired by DeltaVision Elite imaging (GE Healthcare Inc.) software. Tracking C19Q cluster formation was performed for 4 hours and images were captured every 30 minutes. For the representative images, the raw data were deconvoluted by softWoRx (GE Healthcare Inc.) software.

### 3D-SIM procedure

3D-SIM was performed on a DeltaVision OMX V4 system (GE Healthcare) equipped with a 60x (1.42 NA) Plan Apo oil-immersion objective (Olympus) and six laser (405, 445, 488, 514, 568 and 642nm). Image stacks were captured and were reconstructed using softWoRx (GE Healthcare)._Images were registered with alignment parameters obtained from calibration measurements with 100 nm diameter TetraSpeck Microspheres with four colors (Molecular Probes).

### SETDB1 knockdown

SETDB1 siRNA (GGCAAGAAGAGAACUAAGA) and Control siRNA were purchased from GenePharma. EpiGo-KRAB cell lines were transfected with control siRNA or SETDB1 siRNA by SiRNA-Mate (GenePharma) according to the manufacture’s instruction. Cells were collected 48 hours after transfection. For RT-qPCR, total RNAs were isolated using quick-RNA MiniPrep kit (Zymo Research Inc.) according to manufacturer’s instruction. RNA was converted to cDNA using HifairTM II 1st Strand cDNA Synthesis SuperMix (Yeasen). Quantitative PCR was carried out on StepOnePlusTM by using Hieff UNICON^®^ Power qPCR SYBR Green Master Mix (Yeasen). All qPCR data is represented as the mean +/− standard deviation of three replicates. primer sequences for SETDB1 and GAPDH are as follows: SETDB1 forward primer (AGGAACTTCGGCATTTCATCG) and SETDB1 reverse primer (TGTCCCGGTATTGTAGTCCCA); GAPDH forward primer (CGACCACTTTGTCAAGCTCA) and GAPDH reverse primer (AGGGGTCTACATGGCAACTG).

### ChIP-seq

Cells were collected and crosslinked by 1% formaldehyde. Cells were suspended in cell lysis buffer B (50 mM Tris-HCl PH 8.0, 20 mM EDTA, 0.3% SDS, freshly added protease inhibitor) for 10 min on ice before sonication. 1mg beads (Invitrogen) were washed with PBS+5 mg/mL BSA for three times and incubated with 5 μg antibody at 4°C for 6-8 hours before adding chromatin. Sonicated chromatin was diluted in dilution buffer (16.7 mM Tris-HCl PH 8.0, 1.1% Triton X-100, 1.2 mM EDTA, 167 mM NaCl, freshly added protease inhibitor), and was incubate with pre-treated beads and antibodies by rotation at 4°C overnight. Then beads were washed with washing buffer (50 mM Hepes PH 8.0, 1% NP-40, 0.7% DOC, 0.5 M LiCl, freshly added protease inhibitor) for 5 times followed by wash with TE once. 100μL elution buffer (50 mM Tris-HCl pH 8.0, 1 mM EDTA, 1% SDS) was added and beads were incubated in Thermo mixer at 65°C for 30 min with max speed. Supernatant was collected and treated with protease K at 55°C for 2 hours, then purified with DNA purification kit (Tiangen). RNA was removed by treatment of RNase at 37°C for 1 hour. DNA was then purified by AMPure Beads (Beckman) and subjected to DNA library preparation as described below.

### RNA-seq library preparation and sequencing

5 mg RNA was extracted using quick-RNA MiniPrep kit (Zymo) and then treated with DNase I (Fermentas) at 37°C for 1 hour. RNA was then purified using AMPure beads. Poly-A tailed mRNA was collected using Dynabeads™ mRNA purification kit (Invitrogen). Purified RNA was fragmented with RNA Fragmentation Buffer (NEB) at 95°C for 5 min. Reaction was stopped and RNA was purified by AMPure beads. First strand cDNA was synthesized with a commercial kit using both oligo dT and random primers (Invitrogen). Second strand cDNA was synthesized with second strand synthesis buffer (Invitrogen), MgCl_2_, DTT, dNTP, dUTP, RNase H (Fermentas), E. coli DNA ligase (NEB) and DNA polymerase I (NEB). DNA was purified after 2 hours incubation on thermomixer at 16°C. Synthesized cDNA was subjected to DNA library preparation as described below.

### Library construction

Purified DNA or cDNA was subjected to NEBNext Ultra II DNA Library Prep Kit for Illumina (NEB). The DNA was resuspended in 25 μl ddH2O, followed by end-repair/A-tailing with 3.5 μl End Prep buffer and 1.5 μl enzyme mix according to manufactory instruction. The ligation reaction was then performed by adding diluted 1.25 μl adaptors (NEB), 15 μl Ligation master mix, 0.5 μl Ligation enhancer, at 4°C for overnight. The ligation reaction was treated with 1.5 μl USERTM enzyme according to the instruction and was purified by AMPure beads. The 1^st^ round PCR was performed by adding 25 μl 2x KAPA HiFi HotStart Ready Mix (KAPA biosystems) with primers of NEB Oligos kit, with the program of 98°C for 45 s, (98°C for 15 s and 60°C for 10 s) with 8-9 cycles and 72°C for 1 min. DNA was purified using 1x AMPure beads and a 2^nd^ round PCR was performed with 25 μl 2x KAPA HiFi HotStart Ready Mix with Illumina universal primers, and same PCR program for 8 cycles. The final libraries were purified by 1x AMPure beads and subjected to next-generation sequencing. All libraries were sequenced on Illumina HiSeq 2500 or HiSeq XTen according to the manufacturer’s instruction.

### sisHi-C library generation and sequencing

The sisHi-C library generation was performed as described previously [38]. Briefly, spermatogenetic cells were fixed with 1% formaldehyde at room temperature (RT) for 10 min. Formaldehyde was quenched with glycine for 10 min at RT. Cells were washed with 1XPBS for two times and then lysed in 50 μl lysis buffer (10 mM Tris-HCl pH7.4, 10 mM NaCl, 0.1 mM EDTA, 0.5% NP-40 and proteinase inhibitor) on ice for 50 min. After spinning at 3000 rpm/min in 4°C for 5 min, the supernatant was discarded with a pipette carefully. Chromatin was solubilized in 0.5% SDS and incubated at 62°C for 10 min. SDS was quenched by 10% Triton X-100 at 37°C for 30 min. Then the nuclei were digested with 50 U Mbo I at 37°C overnight with rotation. Mbo I was then inactivated at 62°C for 20 minutes. To fill in the biotin to the DNA, dATP, dGTP, dTTP, biotin-14-dCTP and Klenow were added to the solution and the reaction was carried out at 37°C for 1.5 hours with rotation. The fragments were ligated at RT for 6 hours with rotation. This was followed by reversal of crosslink and DNA purification. DNA was sheared to 300-500 bp with Covaris M220. The biotin-labeled DNA was then pulled down with 10ul Dynabeads MyOne Streptavidin C1 (Life Technology). Sequencing library preparation was performed on beads, including end-repair, dATP tailing and adaptor-ligation. DNA was eluted twice by adding 20ul water to the tube and incubation at 66°C for 20 minutes. 9-15 cycles of PCR amplification were performed with Extaq (Takara). Finally, size selection was done with AMPure XP beads and fragments ranging from 200 bp to 1000 bp were selected. All the libraries were sequenced on Illumina HiSeq2500 or HiSeq Xten according to the manufacturer’s instruction.

### Imaging data analysis

The maximum projection of Z stack was used for the quantification of cell numbers and measurement of colocalization analysis of HP1α and H3K9me3 were performed with Fiji software. Analysis trajectory of live cell loci site using MTrack2 Plugins. Fluorescent puncta were identified as local maxima satisfying the minimum intensity and min/max peak with threshold by Gaussian fitting. Data are represented as mean ± SEM. The exact number, n, of data points and the representation of n (cells, independent experiments) are indicated in the respective figure legends and in the Results.

### ChIP data processing

The paired-end ChIP reads were aligned with the parameters: -t –q –N 1 –L 25 –X 1000 --nomixed --no-discordant. All unmapped reads, non-uniquely mapped reads, reads with low mapping quality (MAPQ < 20) and PCR duplicates were removed. For downstream analysis, we normalized the read counts by computing the numbers of reads per kilobase of bin per million of reads sequenced (RPKM) for 100-bp bins of the genome. To minimize the batch and cell type variation, RPKM values across whole genome were further Z-score normalized. To visualize the ChIP signals in the UCSC genome browser, we generated the RPKM values on a 100 bp-window base. H3K9me3 peaks were called using MACS2 [39] with the parameters –g hs –broad – nomodel –nolambda --broad and noisy peaks with very weak signals (RPKM < 5) were removed from further analyses. Adjacent peaks with very close distance (<5kb) were merged for downstream analyses.

### RNA-seq data processing

All RNA-seq data were mapped to hg19 reference genome by Tophat. The gene expression levels were calculated by Cufflinks (version 2.2.1) using the refFlat database from the UCSC genome browser.

### Hi-C data mapping

Paired end raw reads of Hi-C libraries were aligned, processed and iteratively corrected using HiC-Pro (version 2.7.1b) as described [40]. Briefly, sequencing reads were first independently aligned to the human reference genome (hg19) using the bowtie2 end-to-end algorithm and very-sensitive” option. To rescue the chimeric fragments spanning the ligation junction, the ligation site was detected and the 5’ fraction of the reads was aligned back to the reference genome. Unmapped reads, multiple mapped reads and singletons were then discarded. Pairs of aligned reads were then assigned to Mbo I restriction fragments. Read pairs from uncut DNA, self-circle ligation and PCR artifacts were filtered out and the valid read pairs involving two different restriction fragments were used to build the contact matrix. Valid read pairs were then binned at a specific resolution by dividing the genome into bins of equal size. We chose 100-kb bin size for examination of global interaction patterns of the whole chromosome, and 40-kb bin size to show local interactions and to perform TAD calling. Then the binned interaction matrices were normalized using the iterative correction method [40, 41] to correct the biases such as GC content, mappability and effective fragment length in Hi-C data.

### Identification of conventional chromatin compartments and refined-A/B

Conventional chromatin compartments A and B were identified with a method described previously [42] with some modifications. The normalized 100 kb interaction matrices for each stage were used in this analysis. Firstly, the bins that have no interactions with any other bins were removed before expected interaction matrices were calculated. Observed/Expected matrices were generated using a sliding window approach [43] with the bin size of 400 kb and the step size of 100 kb. Principal component analysis was performed on the correlation matrices generated from the observed/expected matrices. The first principal component of the correlation matrices together with gene density were used to identify A/B compartments. In this analysis, the correlation matrices were calculated according to the interaction matrices separated according to the location of predicted centromeres for each chromatin, as the principal component often reflects the partitioning of chromosome arms [42]. As for Refined-A/B compartment, the calling method were similar to the conventional chromatin compartments, with the matrices restricted to 10 Mb one-by-one instead of whole chromosome arms. The PC1 value generated by those restricted matrices were taken as local PC1. Juicebox was used to generate all chromosome-wide as well as the zoom-in views of interaction frequency heatmap in this study. Both the conventional compartment correlation heatmap and the Refined-A/B compartment heatmap were generated with Java TreeView according to the corresponding correlation matrices.

## Additional file

### Supplemental informations

Supplemental Information includes 5 figures, 2 videos, and 3 data files and can be found with this article online.

## Acknowledgements

We thank Luke Lavis (Janelia Research Campus, Howard Hughes Medical Institute, Ashburn, VA, USA) for the HaloTag JF-549. U2OS Genomic DNA was a gift Xingxu Huang. We thank Jianan Li for help with Lentivirus production. We are grateful to Guisheng Zhong, Cuiping Tian, Xiaoming Li and Yi Qian for help with imaging. We thank Pengwei Zhang and Shuangli Zhang for help with cell sorting. OMX4 microscopy was provided by Shanghai Institute for Advanced Immunochemical Studies (SIAIS) at Shanghaitech University and Fluorescence activated cell sorting (FACS) was provided by iHuman Institute of ShanghaiTech. We thank members of the Ma’s laboratory and Xie’s laboratory for their comments during preparation of the manuscript. This work was funded by National Natural Science Foundation of China (No. 31970591 to H. Ma) and the Shanghai Pujiang program (19PJ1408000 to H. Ma).

## Data and code availability

Raw sequence reads of U2OS-EpiGo-Control and U2OS-EpiGo-KRAB experiments are deposited in NCBI GEO: GSE137469. All custom code used in this study is available upon request.

## Authors’ contributions

H.M. conceived this project. H.M. and Y.F. designed the experiments. A.N., X.C. and S.Z. performed data mining of chromosome-specific repeats. C.Y. made the constructs and performed qPCR. X.W. and X.H. performed Oligopaint FISH and ImmunoFISH. Y.F. performed fluorescence imaging and 3D-SIM. Y.W. performed RNA-seq, ChIP-seq and Hi-C. H.M., Y.F., W.X., Y.W., J.Z., X.X., and T.P. interpreted data. Y.F., Y.W., W.X. and H.M. wrote the paper with input from all the authors.

## SUPPLEMENTAL INFORMATION

**Fig. S1.**
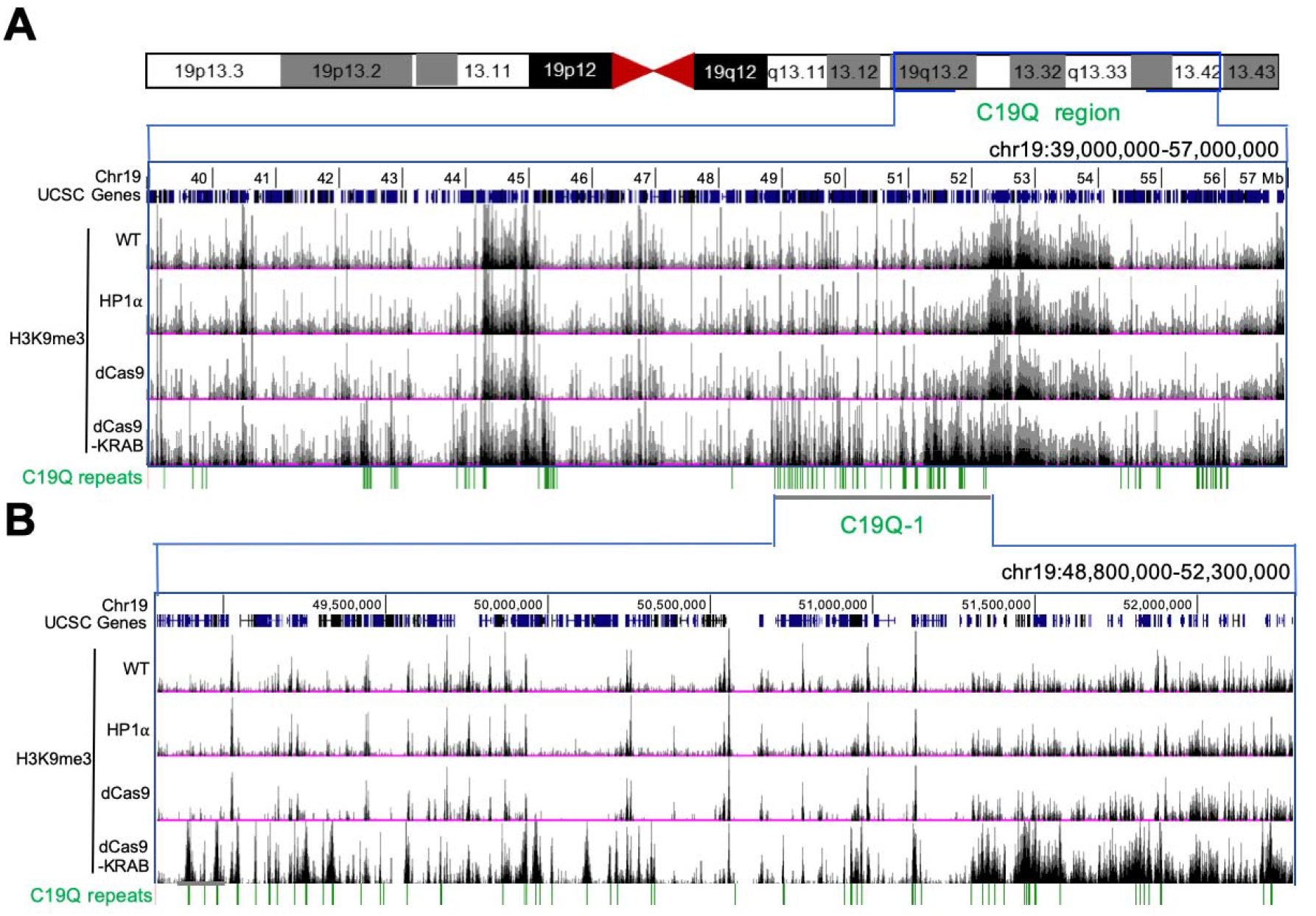
H3K9 trimethylation of C19Q regions mediated by the EpiGo-KRAB system. Related to Figure 1. (A) H3K9me3 states of C19Q region in the cell lines of U2OS wildtype (WT), U2OS-HP1α-HaloTag (HP1α), EpiGo-Control (dCas9) or EpiGo-KRAB (dCas9-KRAB). ChIP-seq of H3K9me3 was performed in these four cell lines. C19Q repeats shown the C19Q target sites. (B) C19Q-1 region is chosen to show the difference of H3K9me3 state between U2OS wildtype (WT), U2OS-HP1α-HaloTag (HP1α), EpiGo-Control (dCas9) or EpiGo-KRAB (dCas9-KRAB) cell lines. C19Q repeats shown the C19Q target sites.

**Fig. S2.**
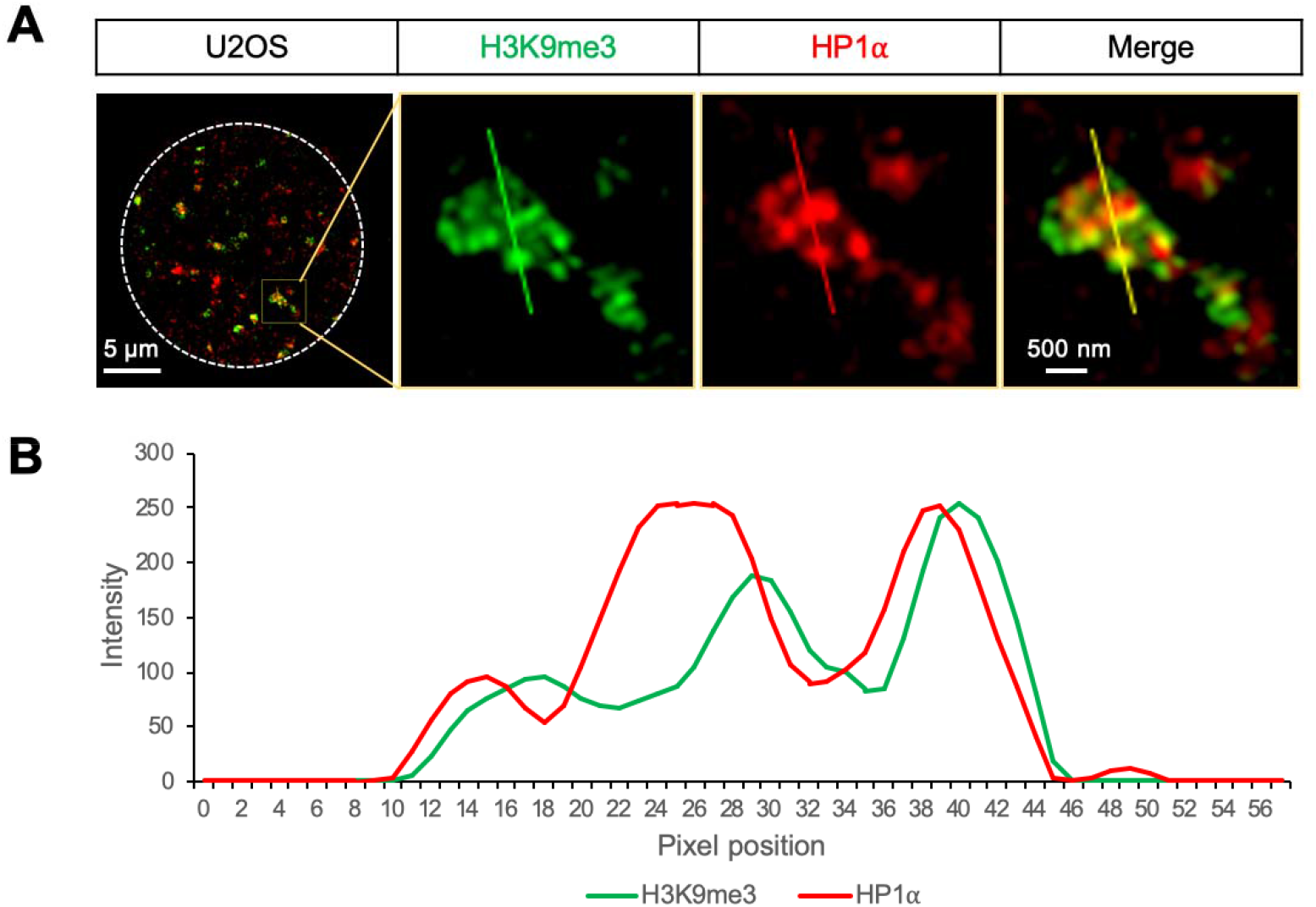
Colocalization of H3K9me3 and HP1α in U2OS cells. Related to Figure 2. (A) Double staining of endogenous H3K9me3 and HP1α. H3K9me3 puncta (green) were detected by H3K9me3 antibody and HP1α (red) was tagged by HaloTag. (B) Linescan of the vertical line. linescan shows, from top to bottom, the intensities (arbitrary units) of H3K9me3 (green) and HP1α (red) along the line indicated in (A).

**Fig. S3.**
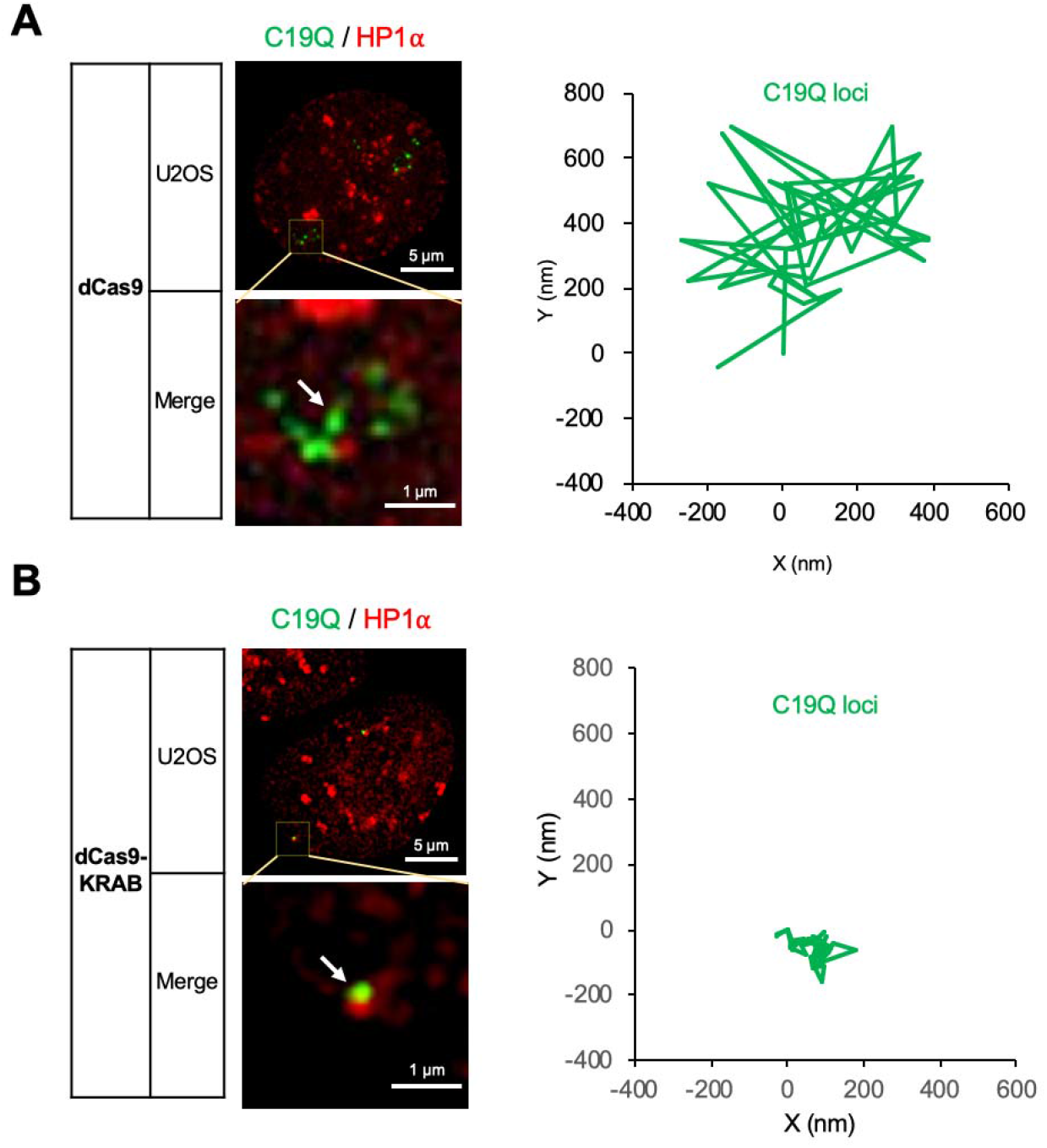
Tracking the Dynamics of C19Q loci in EpiGo-KRAB cells. Related to Figure 2. (A) Live cell tracking of C19Q loci movement in EpiGo-Control cell lines. The dynamics of C19Q (green) and HP1α (red) were tracked for 30 seconds. The movement of locus C19Q loci (arrowed) were recorded at 100 milliseconds per frame for 30 seconds. All trajectories were shifted to start from the origin (0, 0) for easy comparison of the movement vectors. The locus movements were corrected for the possible movements of microscope stage. (B) Live cell tracking of C19Q loci movement in EpiGo-KRAB cell lines. The image processing details are the same as described in (A).

**Fig. S4.**
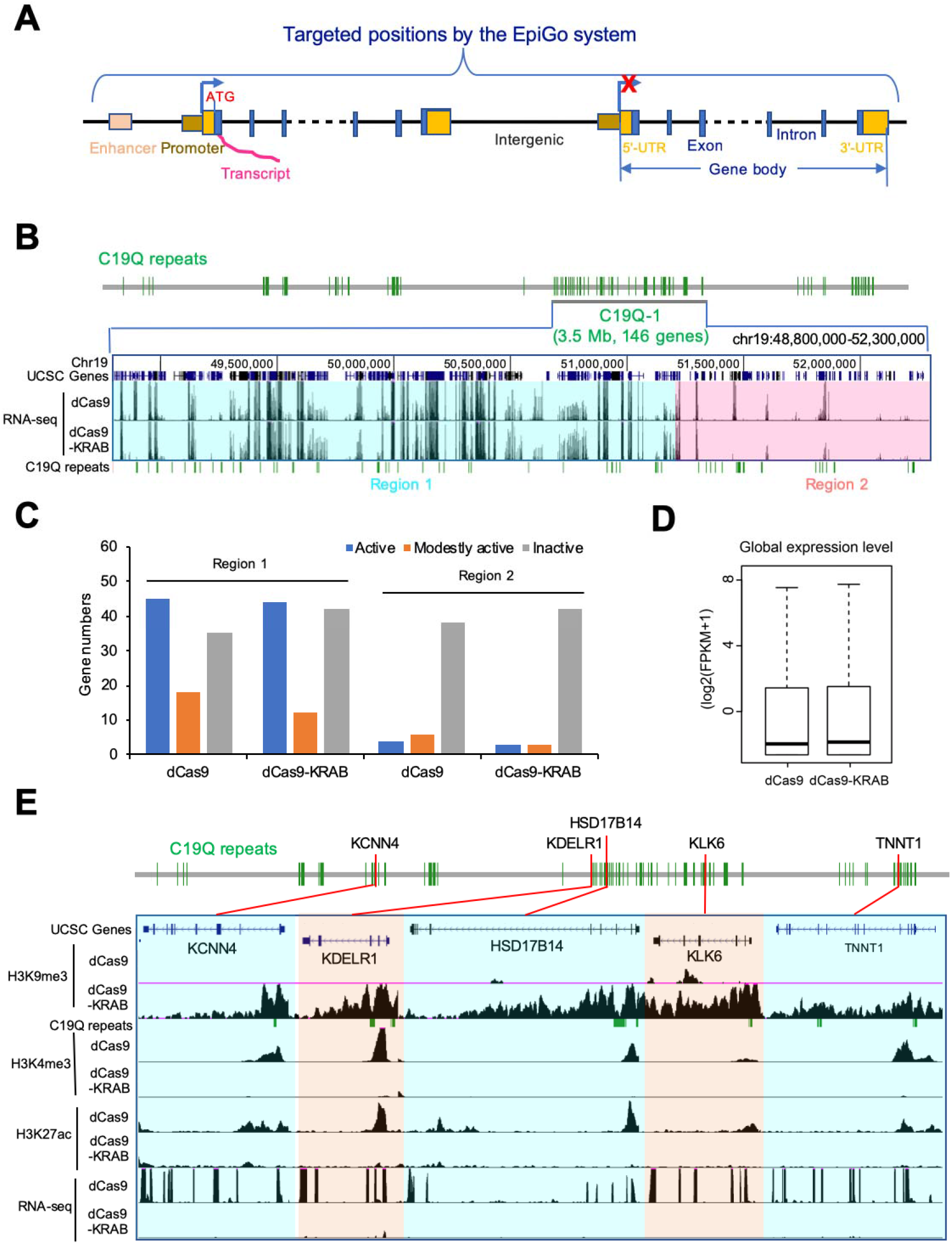
The influence of H3K9me3 on transcription. Related to Figure 3. (A) Diagram of targeted positions by the EpiGo system, which includes enhancer, promoter, 5’-UTR, exon, intron, 3’-UTR and intergenic regions. (B) RNA levels in EpiGo-Control (dCas9) and EpiGo-KRAB (dCas9-KRAB) cell lines. RNA-seq was performed in these two cell lines. Region 1 (chr19: 48,800,000-51,200,000) or Region 2 (chr19:51,200,001-52,300,000) of C19Q-1 with different transcriptional states was highlighted with cyan or pink respectively. (C) Histogram shown active, modestly active and inactive genes in Region 1 and Region 2 in EpiGo-Control (dCas9) and EpiGo-KRAB (dCas9-KRAB) cell lines. Active genes were defined by applying a threshold of FPKM ≥5, modestly active genes were applied a threshold of 1 < FPKM <5 and inactive genes were used a threshold of FPKM ≤1. (D) Global expression level in EpiGo-Control (dCas9) and EpiGo-KRAB (dCas9-KRAB) cell lines. (E) ChIP-seq of H3K9me3, H3K4me3, H3K27ac and RNA-seq were performed in EpiGo-Control (dCas9) and EpiGo-KRAB (dCas9-KRAB) cell lines. *KCNN4, HSD17B14, TNNT1* genes and *KDELR1, KLK6* genes were highlighted with light orange or cyan respectively. All genes were marked location in C19Q repeats with red lines.

**Fig. S5.**
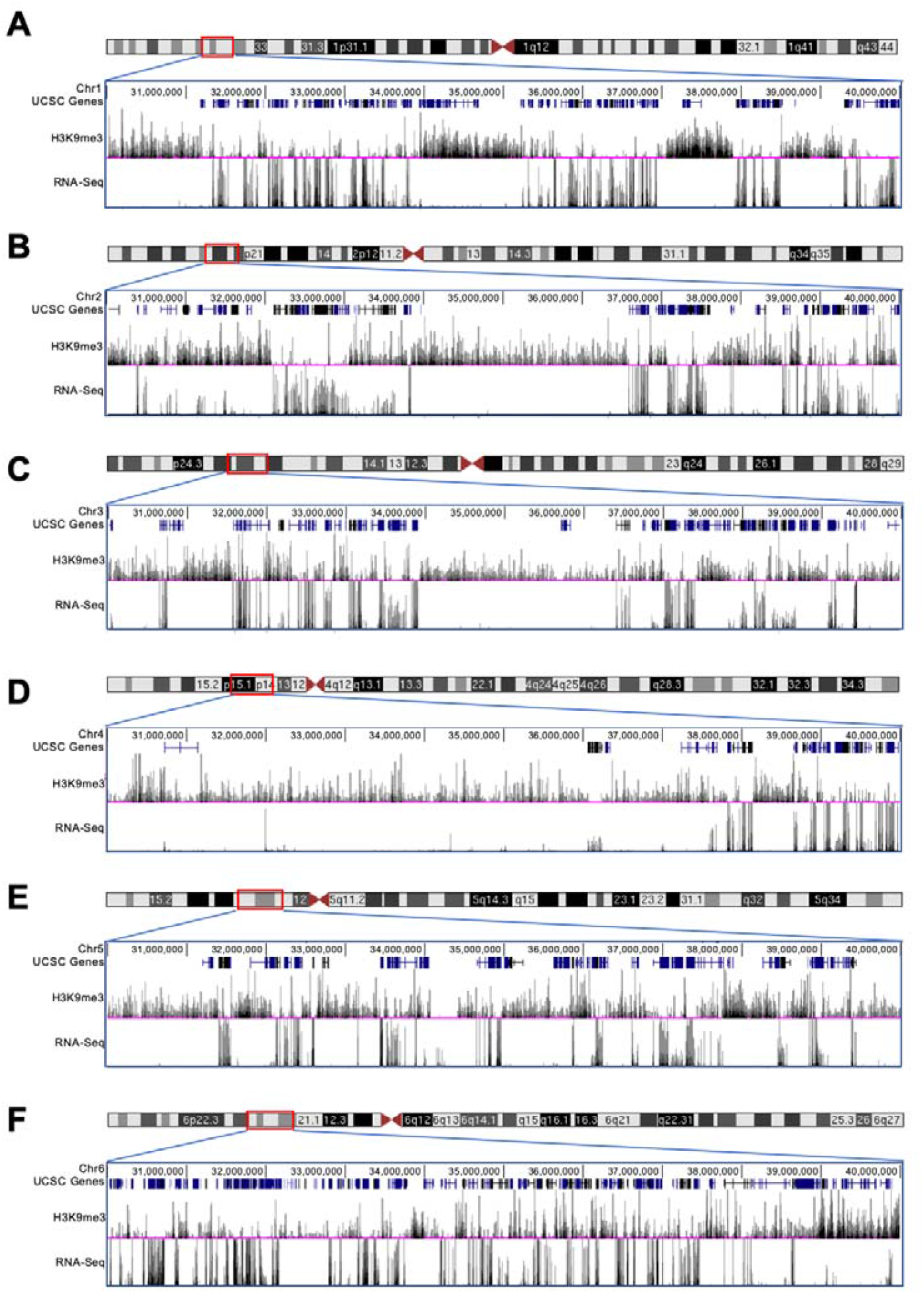
Negative correlation of H3K9 trimethylation and active transcription. Related to Figure 4. The H3K9me3 state and RNA levels of 30-40 Mb regions in chromosome 1-6 were shown in A-F respectively. The ChIP-seq of H3K9me3 and RNA-seq were performed on U2OS-EpiGo-Control cell lines.

## Supplementary Tables

**Table S1. Target sites of C19Q on the q-arm of human chromosome 19.**

**Table S2. A list of regions and primers for Oligopaint FISH probes in This Study.**

**Table S3. A list of template oligonucleotides for Oligopaint FISH probes in This Study.**

## Supplementary Movies

**Movie S1. Tracking the movement of C19Q loci in U2OS-EpiGo-Control cells.**

Images were cropped to 50X50 pixels and each video includes 300 frames (a total time of 30 seconds). The imaging rate is 100 milliseconds per frame and the play rate is 30 frames per second. The individual locus movements were corrected for the possible movements of microscope stage.

**Movie S2. Tracking the movement of C19Q loci in U2OS-EpiGo-KRAB cells.**

The image processing details are the same as described in the Video S1.

## References

1. Dixon JR, Selvaraj S, Yue F, Kim A, Li Y, Shen Y, Hu M, Liu JS, Ren B: Topological domains in mammalian genomes identified by analysis of chromatin interactions. Nature 2012, 485:376–380.

2. de Laat W, Duboule D: Topology of mammalian developmental enhancers and their regulatory landscapes. Nature 2013, 502:499–506.

3. Sima J, Chakraborty A, Dileep V, Michalski M, Klein KN, Holcomb NP, Turner JL, Paulsen MT, Rivera-Mulia JC, Trevilla-Garcia C, et al: Identifying cis Elements for Spatiotemporal Control of Mammalian DNA Replication. Cell 2019, 176:816–830 e818.

4. Gibcus JH, Dekker J: The hierarchy of the 3D genome. Mol Cell 2013, 49:773–782.

5. Nuebler J, Fudenberg G, Imakaev M, Abdennur N, Mirny LA: Chromatin organization by an interplay of loop extrusion and compartmental segregation. Proc Natl Acad Sci US A 2018, 115:E6697–E6706.

6. Stadhouders R, Filion GJ, Graf T: Transcription factors and 3D genome conformation in cell-fate decisions. Nature 2019, 569:345–354.

7. Larson AG, Elnatan D, Keenen MM, Trnka MJ, Johnston JB, Burlingame AL, Agard DA, Redding S, Narlikar GJ: Liquid droplet formation by HP1alpha suggests a role for phase separation in heterochromatin. Nature 2017, 547:236–240.

8. Strom AR, Emelyanov AV, Mir M, Fyodorov DV, Darzacq X, Karpen GH: Phase separation drives heterochromatin domain formation. Nature 2017, 547:241–245.

9. Wang L, Gao Y, Zheng X, Liu C, Dong S, Li R, Zhang G, Wei Y, Qu H, Li Y, et al: Histone Modifications Regulate Chromatin Compartmentalization by Contributing to a Phase Separation Mechanism. Mol Cell 2019, 76:646–659 e646.

10. Falk M, Feodorova Y, Naumova N, Imakaev M, Lajoie BR, Leonhardt H, Joffe B, Dekker J, Fudenberg G, Solovei I, Mirny LA: Heterochromatin drives compartmentalization of inverted and conventional nuclei. Nature 2019, 570:395–399.

11. Rao SS, Huntley MH, Durand NC, Stamenova EK, Bochkov ID, Robinson JT, Sanborn AL, Machol I, Omer AD, Lander ES, Aiden EL: A 3D map of the human genome at kilobase resolution reveals principles of chromatin looping. Cell 2014, 159:1665–1680.

12. Wang S, Su JH, Beliveau BJ, Bintu B, Moffitt JR, Wu CT, Zhuang X: Spatial organization of chromatin domains and compartments in single chromosomes. Science 2016, 353:598–602.

13. Boettiger AN, Bintu B, Moffitt JR, Wang S, Beliveau BJ, Fudenberg G, Imakaev M, Mirny LA, Wu CT, Zhuang X: Super-resolution imaging reveals distinct chromatin folding for different epigenetic states. Nature 2016, 529:418–422.

14. Bintu B, Mateo LJ, Su JH, Sinnott-Armstrong NA, Parker M, Kinrot S, Yamaya K, Boettiger AN, Zhuang X: Super-resolution chromatin tracing reveals domains and cooperative interactions in single cells. Science 2018, 362.

15. Mateo LJ, Murphy SE, Hafner A, Cinquini IS, Walker CA, Boettiger AN: Visualizing DNA folding and RNA in embryos at single-cell resolution. Nature 2019, 568:49–54.

16. Nir G, Farabella I, Perez Estrada C, Ebeling CG, Beliveau BJ, Sasaki HM, Lee SD, Nguyen SC, McCole RB, Chattoraj S, et al: Walking along chromosomes with superresolution imaging, contact maps, and integrative modeling. PLoS Genet 2018, 14:e1007872.

17. Cardozo Gizzi AM, Cattoni DI, Fiche JB, Espinola SM, Gurgo J, Messina O, Houbron C, Ogiyama Y, Papadopoulos GL, Cavalli G, et al: Microscopy-Based Chromosome Conformation Capture Enables Simultaneous Visualization of Genome Organization and Transcription in Intact Organisms. Mol Cell 2019, 74:212–222 e215.

18. Nozaki T, Imai R, Tanbo M, Nagashima R, Tamura S, Tani T, Joti Y, Tomita M, Hibino K, Kanemaki MT, et al: Dynamic Organization of Chromatin Domains Revealed by Super-Resolution Live-Cell Imaging. Mol Cell 2017, 67:282–293 e287.

19. Ma H, Tu LC, Naseri A, Chung YC, Grunwald D, Zhang S, Pederson T: CRISPR-Sirius: RNA scaffolds for signal amplification in genome imaging. Nat Methods 2018, 15:928–931.

20. Chen B, Gilbert LA, Cimini BA, Schnitzbauer J, Zhang W, Li GW, Park J, Blackburn EH, Weissman JS, Qi LS, Huang B: Dynamic imaging of genomic loci in living human cells by an optimized CRISPR/Cas system. Cell 2013, 155:1479–1491.

21. Thakore PI, D’Ippolito AM, Song L, Safi A, Shivakumar NK, Kabadi AM, Reddy TE, Crawford GE, Gersbach CA: Highly specific epigenome editing by CRISPR-Cas9 repressors for silencing of distal regulatory elements. Nat Methods 2015, 12:1143–1149.

22. Klann TS, Black JB, Chellappan M, Safi A, Song L, Hilton IB, Crawford GE, Reddy TE, Gersbach CA: CRISPR-Cas9 epigenome editing enables high-throughput screening for functional regulatory elements in the human genome. Nat Biotechnol 2017, 35:561–568.

23. Ma H, Tu LC, Naseri A, Huisman M, Zhang S, Grunwald D, Pederson T: Multiplexed labeling of genomic loci with dCas9 and engineered sgRNAs using CRISPRainbow. Nat Biotechnol 2016, 34:528–530.

24. Frietze S, O’Geen H, Blahnik KR, Jin VX, Farnham PJ: ZNF274 recruits the histone methyltransferase SETDB1 to the 3’ ends of ZNF genes. PLoS One 2010, 5:e15082.

25. Schultz DC, Ayyanathan K, Negorev D, Maul GG, Rauscher FJ, 3rd: SETDB1: a novel KAP-1-associated histone H3, lysine 9-specific methyltransferase that contributes to HP1-mediated silencing of euchromatic genes by KRAB zinc-finger proteins. Genes Dev 2002, 16:919–932.

26. Groner AC, Meylan S, Ciuffi A, Zangger N, Ambrosini G, Denervaud N, Bucher P, Trono D: KRAB-zinc finger proteins and KAP1 can mediate long-range transcriptional repression through heterochromatin spreading. PLoS Genet 2010, 6:e1000869.

27. Keenen MM, Larson AG, Narlikar GJ, Redding S: Dissecting the Mechanism of HP1 Mediated Chromatin Compaction via Single Molecule DNA Curtains. Biophysical Journal 2018, 114:30a–30a.

28. Leopold K, Stirpe A, Schalch T: Transcriptional gene silencing requires dedicated interaction between HP1 protein Chp2 and chromatin remodeler Mit1. Genes Dev 2019, 33:565–577.

29. Gauchier M, Kan S, Barral A, Sauzet S, Agirre E, Bonnell E, Saksouk N, Barth TK, Ide S, Urbach S, et al: SETDB1-dependent heterochromatin stimulates alternative lengthening of telomeres. Sci Adv 2019, 5:eaav3673.

30. Fortin JP, Hansen KD: Reconstructing A/B compartments as revealed by Hi-C using long-range correlations in epigenetic data. Genome Biol 2015, 16:180.

31. Erdel F, Rademacher A, Vlijm R, Tunnermann J, Frank L, Weinmann R, Schweigert E, Yserentant K, Hummert J, Bauer C, et al: Mouse Heterochromatin Adopts Digital Compaction States without Showing Hallmarks of HP1-Driven Liquid-Liquid Phase Separation. Mol Cell 2020, 78:236–249 e237.

32. Kraushaar DC, Zhao K: The epigenomics of embryonic stem cell differentiation. Int J Biol Sci 2013, 9:1134–1144.

33. Altun G, Loring JF, Laurent LC: DNA methylation in embryonic stem cells. J Cell Biochem 2010, 109:1–6.

34. Miyamoto T, Furusawa C, Kaneko K: Pluripotency, Differentiation, and Reprogramming: A Gene Expression Dynamics Model with Epigenetic Feedback Regulation. PLoS Comput Biol 2015, 11:e1004476.

35. Ma H, Naseri A, Reyes-Gutierrez P, Wolfe SA, Zhang S, Pederson T: Multicolor CRISPR labeling of chromosomal loci in human cells. Proc Natl Acad Sci U S A 2015, 112:3002–3007.

36. Marcais G, Kingsford C: A fast, lock-free approach for efficient parallel counting of occurrences of k-mers. Bioinformatics 2011, 27:764–770.

37. Beliveau BJ, Boettiger AN, Nir G, Bintu B, Yin P, Zhuang X, Wu CT: In Situ Super-Resolution Imaging of Genomic DNA with OligoSTORM and OligoDNA-PAINT. Methods Mol Biol 2017, 1663:231–252.

38. Du Z, Zheng H, Huang B, Ma R, Wu J, Zhang X, He J, Xiang Y, Wang Q, Li Y: Allelic reprogramming of 3D chromatin architecture during early mammalian development. Nature 2017, 547:232.

39. Zhang Y, Liu T, Meyer CA, Eeckhoute J, Johnson DS, Bernstein BE, Nusbaum C, Myers RM, Brown M, Li W, Liu XS: Model-based analysis of ChIP-Seq (MACS). Genome Biol 2008, 9:R137.

40. Servant N, Varoquaux N, Lajoie BR, Viara E, Chen CJ, Vert JP, Heard E, Dekker J, Barillot E: HiC-Pro: an optimized and flexible pipeline for Hi-C data processing. Genome Biol 2015, 16:259.

41. Imakaev M, Fudenberg G, McCord RP, Naumova N, Goloborodko A, Lajoie BR, Dekker J, Mirny LA: Iterative correction of Hi-C data reveals hallmarks of chromosome organization. Nat Methods 2012, 9:999–1003.

42. Lieberman-Aiden E, van Berkum NL, Williams L, Imakaev M, Ragoczy T, Telling A, Amit I, Lajoie BR, Sabo PJ, Dorschner MO, et al: Comprehensive mapping of long-range interactions reveals folding principles of the human genome. Science 2009, 326:289–293.

43. Dixon JR, Jung I, Selvaraj S, Shen Y, Antosiewicz-Bourget JE, Lee AY, Ye Z, Kim A, Rajagopal N, Xie W, et al: Chromatin architecture reorganization during stem cell differentiation. Nature 2015, 518:331–336.

